# Enhancing hypercompact CasΦ2 activity through EPICA.2, an optimized eukaryotic directed evolution platform

**DOI:** 10.64898/2026.08.12.744198

**Authors:** Giulia Vittoria Ruta, Matteo Ciciani, Veronica De Sanctis, Roberto Bertorelli, Chiara Valentini, Daniele Menghini, Eyemen Kheir, Michele Domenico Gentile, Alessio Conci, Antonio Casini, Anna Cereseto

## Abstract

Compact Cas nucleases offer advantages over the widely used SpCas9 due to their smaller size, which enables more efficient delivery for *in vivo* applications. Among these, the phage-encoded CasΦ2 (Cas12j2) is highly promising due to its relaxed PAM requirement (5’-TTN-3’) and compact size (757 aa); however, its translational potential is limited by low editing activity. To enhance the efficacy of CasΦ2, we optimized the previously reported EPICA system, developing EPICA.2, a eukaryotic directed evolution platform to improve nucleases with nearly undetectable activity. EPICA.2 integrates additional yeast evolution rounds to enrich for active variants along with a low background mammalian reporter system that improves detection and selection of enhanced variants. Finally, we set up a long-read sequencing protocol which uses unique molecular identifiers (UMIs) to reduce sequencing errors, enabling accurate identification of the mutation combinations in each evolved variant. Among the most frequent variants, we obtained evoCasΦ2, which contains six activity-boosting mutations with a synergistic effect not predictable by rational engineering. Overall, evoCasΦ2 showed up to 70-fold increased activity in human cells compared to wild-type and outperformed variants generated through rational approaches, highlighting the potential of EPICA.2 as a powerful strategy to evolve genome editing tools with low native activity.

## INTRODUCTION

The efficacy of clustered Regularly Interspaced Short Palindromic Repeats (CRISPR)-CRISPR associated (Cas) systems for genome editing enabled the development of numerous advanced therapies for genetic diseases [1]. Despite these achievements, several challenges still limit a wider use of CRISPR therapies in the clinic. In particular, the molecular size of the most commonly used *Streptococcus pyogenes* Cas9 (SpCas9) [2] nuclease poses challenges for its delivery to target cells *in vivo*, especially when relying on adeno-associated virus (AAV) vectors, which remain the most commonly used platform for gene therapy [3]. Smaller RNA-guided nucleases have been identified, including compact Cas9 and Cas12 variants [4–8] and ancestral insertion sequences Cas9-like OrfB (IscB) and transposon-associated Protein B (TnpB) proteins [9,10]. While these proteins are typically encoded by prokaryotes, specific clades have also been identified in bacteriophages and eukaryotes [11–13]. Notably, the CasФ clade (Cas12j) is exclusively found in huge bacteriophages and includes the hypercompact Cas12 ortholog CasФ2 [11]. Despite its compact size (757 amino acids) and broad protospacer adjacent motif (PAM) compatibility (5’-TTN-3’ PAM), which make it an attractive candidate for human applications, its low editing efficiency in human cells limits its translational potential [11,14].

Various methods have been applied to increase the activity of CRISPR-Cas nucleases, including structure-guided rational engineering, directed evolution and artificial intelligence (AI)-based approaches [15–17]. To steer Cas nuclease function, we previously developed a Eukaryotic Platform to Improve Cas Activity (EPICA) through directed evolution [18]. This approach includes a yeast-based selection step for activity enhancement, building on a previously established strategy to improve fidelity [19], followed by additional screening of highly active mutants in human cells. Here we evolved CasΦ2, a nuclease with extremely limited editing efficiency, using the EPICA platform, which we further optimized to enable the enhancement of Cas orthologs with very low intrinsic activity,developing of EPICA.2. The platform implementation included additional selection cycles in yeast without mutagenesis, enabling the enrichment and amplification of functional variants. In addition, the screening in human cells was performed using a Cas-induced frameshift reporter system with minimal background signal, allowing the selection of variants with significantly improved activity while excluding poorly performing ones. Finally, since one of the major hurdles of directed evolution experiments is the reconstruction of the exact combination of mutations characterizing each evolved variant [20], we adapted a long-read sequencing approach with error correction to identify full-length variants [21]. Using this improved platform we generated evoCasФ2, a variant with enhanced activity relative to both wild-type and previously engineered variants [14], demonstrating the power of this approach for Cas nucleases optimization.

## RESULTS

### Enhancement of CasΦ2 activity through EPICA.2, an optimized eukaryotic directed evolution platform

To evolve CasΦ2, the EPICA platform [18] was adapted for the evolution of nucleases with low activity. As previously described [18], a *Saccharomyces cerevisiae* strain was engineered by integrating two cassettes in the tryptophan 1 (TRP1) and adenine 2 (ADE2) loci, each containing distinct CasΦ2 PAM and target sequences flanked by homologous arms (**Fig. 1a**). With this setup, the CasΦ2-mediated cleavage triggers single-strand annealing (SSA) repair at the homologous arms, which restores TRP1 and ADE2 function, enabling the selective survival and isolation of active variants on TRP/ADE deficient plates. The activity of CasΦ2 in yeast was evaluated by comparing it with the gold standard SpCas9: while SpCas9 yielded very high colony survival (∼100%) in all the three conditions (TRP–, ADE– and TRP–ADE–), CasΦ2 showed only minimal survival (∼5%) by targeting the ADE2 cassette (**Fig. 1b**). Instead, targeting the TRP1 cassette alone, or in combination with ADE2, CasΦ2 yielded almost no colonies, indicating a stronger stringency by TRP–selection.

**Figure 1.**
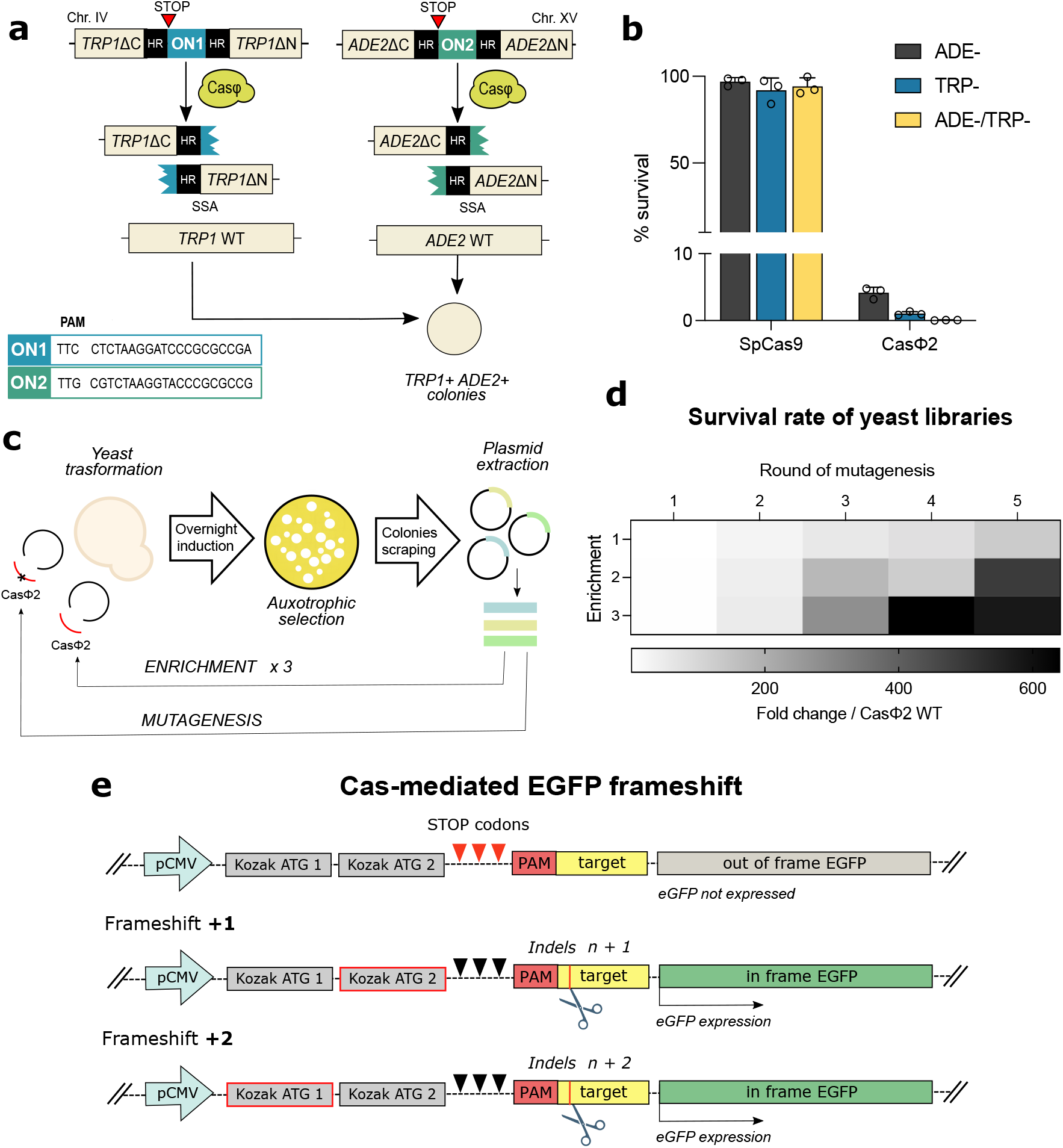
Enhancement of CasΦ2 activity through EPICA.2. **a**) Schematic representation of the yeast strain for CasΦ2 enhancement; framed sequences correspond to the ON-target 1 (ON1) and ON-target 2 (ON2) cassettes carrying the CasΦ2 PAM and target customized sites. Survival of active variants in ADE−TRP− plates is obtained upon CasΦ2 cleavage on the integrated cassettes and repair by SSA at the homologous arms, with the restoration of ADE2 and TRP1 genes. **b**) Comparison of SpCas9 and CasΦ2 cleavage activity in yeast expressed as percentage of survival, corresponding to the number of colonies growing on TRP−, ADE- and TRP-ADE− plates relative to the total number of transformants measured under unrestricted growth conditions. **c**) Schematic overview of the experimental workflow for the CasΦ2 evolution for activity in yeast; active mutants cleaving the ADE2 and TRP1 cassettes are amplified from colonies growing under auxotrophic selection and undergo additional rounds of mutagenesis and selection. At each round of mutagenesis, three rounds of enrichment are performed by transforming yeast with variants amplified without introducing new mutations. **d**) Heatmap of survival rates of CasΦ2 libraries expressed as fold-change of yeast survival under TRP− or TRP-ADE− selection relative to CasΦ2 WT. Extended data are shown in **Figure S1a-e**. **e**) Scheme of the Cas-mediated EGFP frameshift reporter cell line used for the screening of CasΦ2 variants in human cells. EGFP in the reporter is not expressed (grey); when CasΦ2 induces indel formation on PAM (red) + target (yellow) customized sequence, cleavage events generating +1 or +2 frameshifts place the downstream EGFP coding sequence in the same frame of, respectively, the second or the first Kozak ATG sequence, determining EGFP expression (green). Validation of the reporter cell line is shown in **Figure S2a-b**.

Given the minimal activity of CasΦ2, we had to establish conditions enabling the initiation of the evolution campaign. The first evolution round was performed using a single selection site (TRP1) and a low mutagenesis rate to minimize the probability to introduce detrimental mutations along with beneficial ones (**Fig. S1a**). Moreover, to enrich the newly generated variants, each mutagenesis step was followed by three transformation cycles to favor the accumulation of highly enhanced mutants over poorly active ones (enrichment cycles) (**Fig. 1c**). The double-cleavage selection (TRP–ADE–) was introduced in the second round of evolution, following the emergence of improved variants obtained from the first cycle. We carried out successive rounds of mutagenesis combined with enrichment cycles until survival began to decline due to the accumulation of deleterious mutations (**Fig. 1d, Fig. S1b-e**). The top-performing library emerged in the third enrichment of the fourth evolution round achieving a 640-fold increase in yeast survival compared to CasΦ2 wild-type (WT); these mutants were chosen for further selection in human cells. To enrich for highly enhanced variants, we developed a stringent reporter system based on Cas-mediated restoration of an out of frame fluorescent cassette (**Fig. 1e**). The reporter construct contains an enhanced green fluorescent protein (EGFP) cassette downstream of the CasΦ2 target site that is not expressed as it is out of frame with two alternative ATG start codons and Kozak sequences. Following the generation of frameshifts by CasΦ2-mediated indels (insertions or deletions), the EGFP coding sequence is reframed with one of the two upstream Kozak-ATG sequences, resulting in activation of green fluorescence. This system enables detection of EGFP expression obtained by two out of three possible frameshifts generated by CasΦ2 mediated cleavages. The reporter cell line exhibited 0% background fluorescence and high stringency, as shown by the low percentage of EGFP-positive cells generated upon SpCas9 cleavage (∼6%) and no positivity in the absence of SpCas9 expression (**Fig. S2a**). The correlation between SpCas9-mediated EGFP activation and indel formation at the target site was confirmed by Tracking of Indels by DEcomposition (TIDE) analysis (10% indels, **Fig. S2b**).

Following four rounds of evolution in yeast, we generated a lentiviral library of variants that were transduced in the reporter cell line and sorted for EGFP-positive cells. The CasΦ2 variant library showed significantly higher EGFP activation compared to CasΦ2 WT, demonstrating the presence of CasΦ2 mutants with enhanced activity (**Fig. S2c**). To confirm the success of the screening process through cell sorting, we quantified indels at the reporter target site in the isolated cells, detecting ∼70% of indels through Sanger sequencing (TIDE analysis). These results indicated the successful selection of CasΦ2 variants with enhanced activity compared to the native enzyme, following yeast evolution rounds and screening with the stringent out-of-frame EGFP reporter system.

### Identification of evolved CasФ2 variants through error-corrected nanopore sequencing

To accurately reconstruct the CasΦ2 variants generated through directed evolution, we adapted an Oxford Nanopore sequencing protocol that incorporates unique molecular identifiers (UMIs) to enable error correction [21] as well as precise quantification of the frequency of each variant in the sequenced library. Two initial rounds of PCR were used to introduce UMI pairs at the 5’ and 3’ of each variant (**Fig. S3a**), which were then amplified using external primers (**Fig. S3b**) and sequenced on the MinION platform (**Fig. S3c**). After sequencing, >1.2 million raw reads were separated into 44,874 distinct UMI bins and error-corrected consensus reads were generated for each bin. A total of 4,130 CasФ2 variants were reconstructed, each carrying unique combinations of non-synonymous substitutions. Variant frequencies in the library were quantified from the number of distinct UMI bins corresponding to each unique variant (**Fig. S3c-e**).

Clustering variants according to shared substitutions revealed multiple clusters harboring distinct combinations of mutations (**Fig. 2a**), reflecting divergent evolutionary trajectories. Analysis of the cumulative mutation distribution along the protein length enabled the identification of six high frequency substitutions (E9K, K29R, G138R, E168K, T355A and I585V), with E168K and T355A present in nearly all variants (>99.7%). To select the best candidate for further characterization, variants were ranked by their frequency in the sequenced library (**Fig. 2b**). The eight most common variants presented different combinations of the six most frequent substitutions, with the most common variant (∼3% of UMI-corrected reads) harboring all of them (**Fig. 2c**).

**Figure 2.**
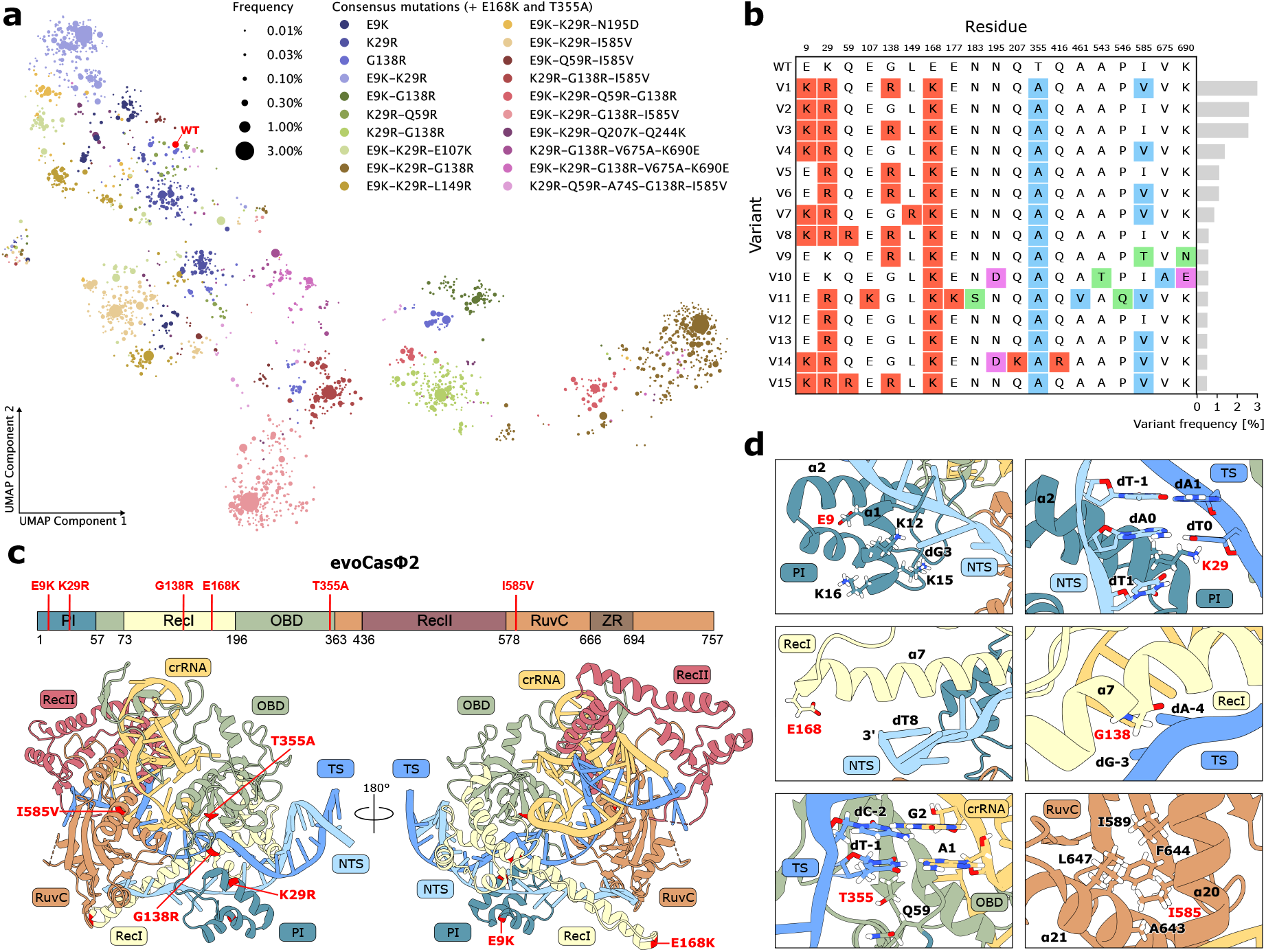
Long-read sequencing of CasФ-2 variants. **a**) UMAP visualization of 2,382 CasФ2 variants reconstructed by long-read sequencing more than once. Variants are clustered by K-means using non-synonymous mutations one-hot encodings. All variants share mutations E168K and T355A, while additional consensus mutations for each cluster are reported. The wild-type variant (not observed in sequencing) is plotted in red as reference. **b**) Most common haplotypes reconstructed by long-read sequencing. Haplotype frequency in the library is reported on the left. Amino acid substitutions are colored according to physiochemical properties: positively charged (red), negatively charged (purple), polar (green), hydrophobic (blue). Wild-type residues and positions are reported in the first row. **c**) Position of the 6 residues mutated in evoCasФ2 highlighted in the wild-type ternary complex (PDB 7LYS). **d**) Close-up view of each mutated residue in the ternary complex.

The most frequent variant was named evoCasФ2. To gain insight into how the six mutations may affect the cleavage activity, the substitutions were mapped onto the CasФ2 WT ternary complex structure [14] (**Fig. 2c-d**). E9 is located in the *a*1 helix of the PAM-interacting (PI) domain, on the solvent-facing side, which is rich in positively charged residues (K12, K15 and K16). E9K increases the positive charge of this helix, possibly favouring target DNA unwinding. K29 is also located in the PI domain, in the *a*2 helix, inserted into the major groove at the 3’ end of the PAM. It was previously shown that K29A impairs DNA binding ability [14], suggesting that the K29R substitution could improve DNA unwinding and R-loop formation by stabilizing the interaction with the PAM. G138 is located in the *a*7 helix of RecI, close to the PAM-proximal target strand backbone. G138R likely improves R-loop formation by interacting with the negatively-charged sugar phosphate backbone. E168 is positioned in the negatively charged tip of the *a*7 helix of RecI, which regulates accessibility to the RuvC active site [14]. It was previously shown that replacing this region with a GSSG linker or introducing mutations which remove the negative charge (including E168A) results in improved cleavage kinetics of both the target and non-target strands [14]. Therefore, E168K likely has a similar effect by replacing the negative charge with a positive one. T355 is located in the oligonucleotide-binding (OBD) domain, at the base of the target-strand:spacer heteroduplex, at the PAM proximal side. The effect of T355A is difficult to interpret, however given its position it may contribute to heteroduplex stabilization. I585 is located in the *a*20 helix of the RuvC domain, at the interface with *a*21, forming a hydrophobic core with I589, A643, F644 and L647. I585V may influence the stability of this core, possibly reducing steric hindrance or facilitating conformational changes.

### EvoCasΦ2, a CasΦ2 variant with enhanced editing efficacy

To evaluate the genome editing efficacy of evoCasΦ2, we compared indel formation with that of CasΦ2 WT in HEK293T cells (**Fig. 3a**). Following previous reports [30], we optimized the native CasΦ2 precursor CRISPR RNA (pre-crRNA) sequence and spacer length to further enhance activity (**Fig. S4a**). EvoCasΦ2 displayed an 8-fold increase in activity relative to CasΦ2 WT at the benchmark locus HEK site 2 (**Fig. 3a**, **Fig. S4b**). We then verified whether any of the six mutations in evoCasΦ2 individually drove its activity; none of the single mutants recapitulated the enhanced activity of evoCasΦ2 and instead exhibited activity levels similar to CasΦ2 WT (**Fig. 3a**). Therefore, the combination of amino acid substitutions obtained through directed evolution has a sinergic effect on the enhancement of activity. To further evaluate if all six mutations were required, we generated variants lacking each mutation individually. Each removal slightly reduced the activity, confirming that all mutations contribute to maximal performance (**Fig. 3b**).

**Figure 3.**
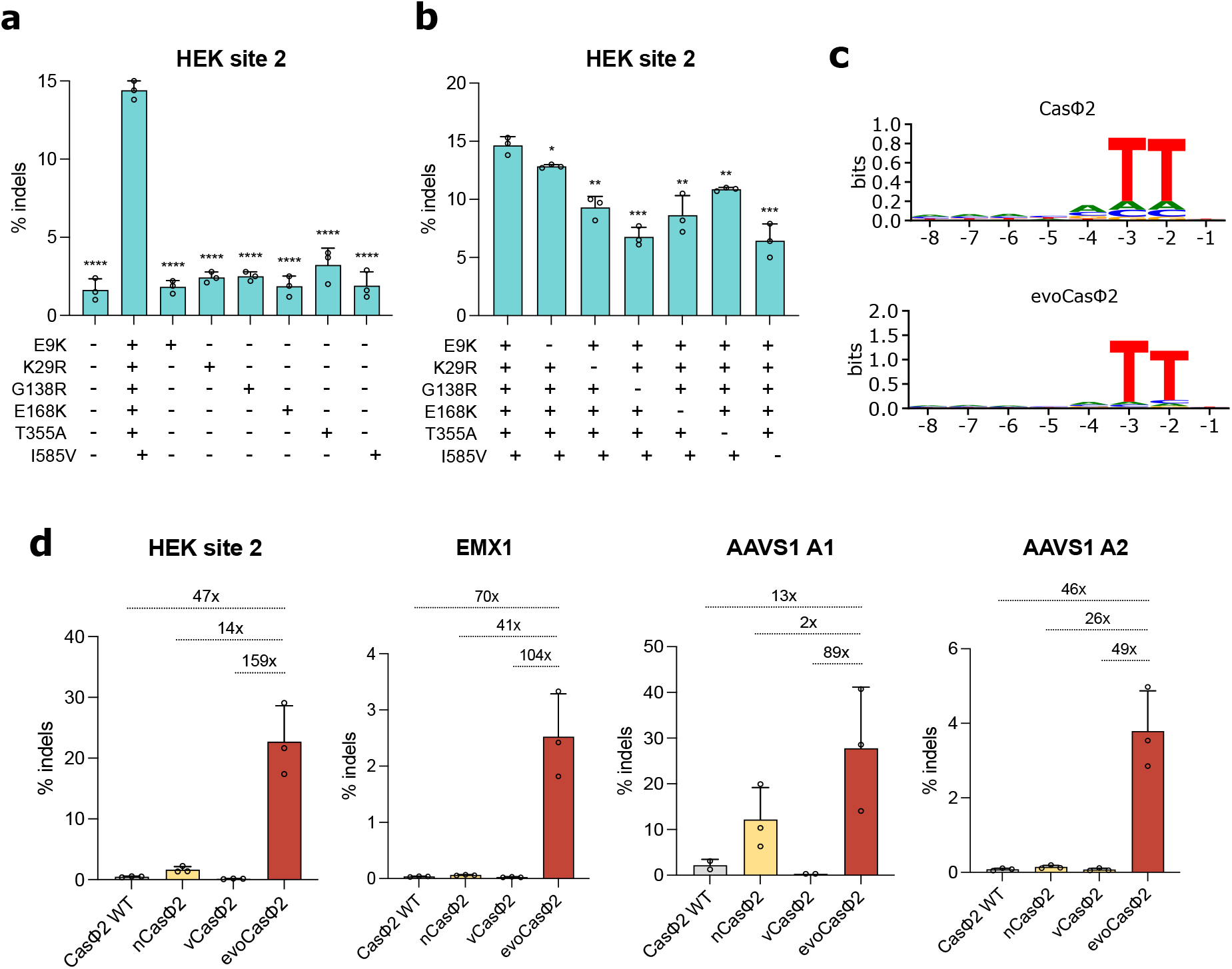
Characterization of evoCasΦ2. **a**) Editing activity at the HEK site 2 locus of CasΦ2 WT, evoCasΦ2 and mutants containing individual amino acid substitutions measured by TIDE analysis in HEK293T cells after plasmids transfection. Plus and minus symbols represent respectively the presence or the absence of individual amino acid substitutions in each variant. **b**) Editing activity of evoCasΦ2 and mutants containing multiple amino acid substitutions measured by TIDE at the HEK site 2 locus in HEK293T cells. **c**) Sequence logo representation of CasΦ2 WT and evoCasΦ2 PAM preferences determined by *in vitro* PAM assay. Heatmaps representing each nucleotide combination are reported in **Figure S5**. **d**) Editing efficiency of CasΦ2 WT, nCasΦ2, vCasΦ2 and evoCasΦ2 in endogenous genomic loci of HEK293T cells; cells were analyzed by deep amplicon sequencing after plasmids transfection. Extended data are reported in **Supplementary Table S1**. In **a**, **b** and **d** data are reported as mean ± standard deviation of n ≥ 2 biologically independent samples. Individual values are represented as empty circles. Statistical analysis in **a** and **b** were performed using unpaired t test: * P < 0.1, ** P < 0.01, *** P < 0.001, **** P < 0.0001.

Since two evoCasΦ2 mutations (E9K, K29R) lie within the PI domain of the protein, we assessed whether the PAM recognition sequence was modified. Results from an *in vitro* PAM-screening assay showed that evoCasΦ2 retained the native CasΦ2 PAM preference, 5’-TTN-3’ [14] (**Fig. 3c**).

Finally, we compared the editing efficacy of evoCasΦ2 to nCasΦ2 and vCasΦ2, two previously reported variants obtained through rational engineering by introducing positively charged residues near the RNA-DNA heteroduplex to increase affinity to the DNA [14,31]. The comparative analysis performed across four genomic loci (HEK site 2, EMX1, AAVS1-A1, AAVS1-A2) showed that evoCasΦ2 has significantly higher editing efficiency at all sites, achieving up to a 70-fold improvement over wild-type CasΦ2 and up to 41-fold and 159-fold higher activity than nCasΦ2 and vCasΦ2, respectively (**Fig. 3d**).

## DISCUSSION

CRISPR-Cas systems have transformed genome editing, emerging as foundational tools across biology and biomedicine [1]. Their remarkable diversity in structure and properties empowers researchers to select nucleases better suited for specific experimental and therapeutic applications [32–34]. A major limitation to the broader exploitation of CRISPR-Cas systems for genome editing applications in eukaryotes is the reduced activity of these enzymes when they are transferred from prokaryotic to eukaryotic environments [35,36]. CasΦ2, a phage-derived nuclease, represents a promising enzyme due to its compact size, which increases its compatibility with cellular delivery systems, and its minimal PAM requirement, which broadens genomic targeting range [14]. Nonetheless, the use of CasΦ2 is limited by a very low editing efficacy in mammalian cells.

Here, we developed EPICA.2 a directed evolution platform which generated evoCasΦ2, a variant capable of measurable editing in human cells, starting from a native enzyme with minimal activity at mammalian genomic sites. At specific sites evoCasΦ2 exhibited low absolute activity (below 5%), yet still produced substantial fold improvements (up to 70 folds) relative to CasΦ2 WT.

Comparative analysis extended to the rationally designed CasΦ2 variants [14] revealed the superior performance of evoCasΦ2. While nCasΦ2 showed modest gains of activity over CasΦ2 WT, vCasΦ2 unexpectedly displayed reduced efficiency despite incorporating positively charged substitutions intended to strengthen DNA interactions. This result can be anyway explained by the fact that neither rationally engineered variant had been previously evaluated for activity in human cells. Notably, evoCasΦ2 shares a single mutated residue with the engineered variant nCasΦ2 (E168K in evoCasΦ2 and E168A in nCasΦ2), yet this single substitution is not sufficient to enhance the editing activity. This observation highlights the strength of an unbiased evolutionary approach over rational design strategies [18,19,37,38] as it can reveal beneficial modifications that are difficult to predict. Indeed, while the effect of the G138R and E168K mutations may be explained by their positive charges in proximity to the DNA-RNA duplex, the functional contributions of the remaining four amino acid substitutions in evoCasΦ2 are not readily predictable.

To precisely identify the combination of mutations present in each variant generated by directed evolution, we adapted an accurate long-read sequencing workflow [21] that also allowed us to quantify the frequency of each variant after selection. We demonstrated that the combination of six mutations in evoCasΦ2 outperformed all individual substitutions, exhibiting a synergistic enhancement not predictable by rational approaches. Long-read sequencing was key to resolve mutation pairings and higher-order combinations that would have been challenging to identify by screening single substitutions or by short-read sequencing approaches. Furthermore, each of the six mutations contributes positively to the activity of evoCasΦ2 at the HEK site 2 locus, highlighting the critical role of long-read sequencing in identifying amino acid substitutions that enhance enzymatic activity.

Importantly, the mammalian reporter system, which exhibits zero background fluorescence, can be customized with user-defined PAM and target sequences, enabling potential selection of variants tailored to specific genomic sites. Furthermore, varying the PAM and target sequences allows the selection stringency to be adjusted according to the nuclease’s native activity on the chosen sequence.

EvoCasΦ2 represents a promising tool for genome editing, particularly due to its minimal PAM sequence (5′-TTN-3′), which enables broad target accessibility, and its compact size, compatible with delivery via AAV vectors; further studies will be required to evaluate its efficiency, specificity and immunogenicity for potential *in vivo* applications.

More broadly, EPICA.2 could be applied to additional nucleases, including the promising compact TnpB and IscB families, to uncover key residues that boost activity in human cells and could serve as a powerful tool for the next generation of genome editing technologies.

## MATERIALS AND METHODS

### Plasmids

For the generation of the yeast strain, the constructs were generated by cloning customized CasΦ2 target and PAM sequences as annealed oligonucleotides into the pUC19 plasmid containing ADE and TRP cassettes, following digestion with KpnI and BamHI. For yeast expression, the CasΦ2 coding sequence was amplified from the pPP441 (Addgene 158801) and cloned in the plasmid p415-GalL-Cas9-CYC1t (Addgene 43804) carrying LEU2, by double digestion with SpeI/XhoI. For sgRNA expression in yeast, pSNR52-BsmBI-CasΦ2_sgRNA was generated cloning CasΦ2 pre-crRNA, BsmBI sites and SUP4 terminator as annealed oligonucleotides in the pSNR52-BsmBI-Cj_sgRNA previously described [18], digested with BsmBI and SpeI. Yeast plasmids carrying CasΦ2 sgRNAs targeting the ON1 and ON2 sites were generated by cloning annealed oligonucleotides into the pSNR52-BsmBI-CasΦ2_sgRNA previously digested with BsmBI. The fragments SNR ON1 and ON2 CasΦ2 sgRNA were then cloned in the pRS316 plasmid carrying URA3 by double digestion with XhoI/SacII. For the sgRNA double plasmid, the pRS316-SNR52p-ON1 CasΦ2 gRNA was digested with SacI and used as backbone to clone the amplified SNR − ON2 CasΦ2 sgRNA fragment. To obtain plasmids carrying SpCas9 sgRNAs, ON1 and ON2 target sequences were introduced in the p426-SNR52p-gRNA.CAN1.Y-SUP4t (Addgene 43803) through PCR site-directed mutagenesis. For the generation of the EGFP out of frame 293 reporter cell line, a DNA sequence containing the two ATG Kozak sequences and the CasΦ2 customized PAM and target was added upstream EGFP sequence with PCR amplification; the DNA fragment was cloned in the pcDNA5 by double digestion with HindIII and EcoRV to obtain the final construct for cell transformation. For expression in mammalian cells, BbsI sites were cloned as annealed oligos in the pPP441 in place of SapI sites; the U6 CasΦ2 sgRNA BbsI fragment was amplified by the pPP441 BbsI previously generated and cloned in the pUC19-SpCas9-opt-sgRNA digested with EcoRI/SalI in place of SpCas9 sgRNA construct. Hepatitis delta virus (HDV) ribozyme was then added to the sgRNA construct by cloning of annealed oligonucleotides in the pUC19-CasΦ2-sgRNA-BbsI digested with BbsI and XbaI; pre-crRNA optimization was obtained modifying the 4 nucleotides CACG in TACG (**Fig. S4a**) by PCR site-directed mutagenesis. Target sequences were cloned as phosphorylated annealed oligonucleotides in the pUC19-CasΦ2-opt-sgRNA-BbsI-HDV previously digested with BbsI and dephosphorylated. For mammalian cell expression, the CasΦ2 coding sequence was PCR-amplified from plasmid pPP441 adding nuclear localization signals (NLS) at both termini as previously described [22] and double digested with AgeI and NheI. The resulting fragment was then cloned into the NcoI/SacI–digested pX330 vector, placing it downstream of the CMV promoter and upstream of puromycin resistance (Addgene Plasmid #194967). EvoCasΦ2, single mutants, multiple mutants, nCasΦ2 and vCasΦ2 were obtained through PCR site-directed mutagenesis performed on pX-CMV-CasΦ2-puro plasmid. For PAM *in vitro* determination assay, CasΦ2 and evoCasΦ2 were PCR-amplified and cloned in the pT7CFE1-NHis-GST-CHA by double digestion with EcoRI and NotI. Primers used for plasmid cloning are reported in **Supplementary Table S2**.

### Yeast culture

The TRP1 ON1 – ADE2 ON2 yeast strain was cultured in YPDA rich medium. For auxotrophic selection, synthetic minimal medium (SD) was used for yeast growth excluding single amino acids on experimental basis; prior to transformations, the yeast strain stably expressing CasΦ2 sgRNA was maintained in SD medium without uracil. Protein expression of CasΦ2 was induced by replacing dextrose in the SD medium with 20 g/L D-( +)-galactose and 10 g/L D-( +)-raffinose.

### Cell culture

HEK293T obtained from the American Type Culture Collection (ATCC) were maintained in Dulbecco’s modified Eagle’s medium (DMEM; Life Technologies) supplemented with 10% FCS (Life Technologies) and antibiotics (Life Technologies). The EGFP OOF (out of frame) 293 cell line was generated by modifying the Flp-In™ 293 Cell Line (Invitrogen) for the expression of the out of frame EGFP construct according to the manufacturer’s protocol. Cell lines were confirmed to be free of mycoplasma contamination (PlasmoTest, Invitrogen).

### Yeast screening

The yeast strain used for the screening was derived from yLFM-ICORE using the Delitto Perfetto approach and the constructs amplified from pUC19-Ade2-ON2 and pUC19-Trp1-ON1. The CasΦ2 variant library was generated through random mutagenesis of the wild-type sequence in the p415-GalL-CasΦ2-CYC1t by error-prone PCR (GeneMorph II kit, Agilent) (primers listed in **Supplementary Table S2**). Mutagenesis conditions were set adjusting the manufacturer’s guidelines to achieve a rate of 0–2 mutations/kb; mutation rate was confirmed by cloning the epPCR in a pcDNA3 vector and Sanger sequencing random colonies to estimate the average number of mutations for kilobases. Yeast cells were co-transformed with the PCR library and the SpeI/XhoI-digested p415-GalL-CYC1t plasmid at a 3:1 ratio to enable *in vivo* assembly. Transformations were carried out following the protocol previously described [23]. After transformation, yeast cells were incubated for 5 hours in SD medium lacking uracil and leucine to allow recovery and recombination, after which CasΦ2 expression was induced by overnight growth in galactose-containing medium. The following day, cells were plated on medium lacking tryptophan or adenine and tryptophan to select colonies harboring improved efficiency variants. The selected colonies were harvested, and plasmids carrying the mutants were isolated using the Zymoprep Yeast Plasmid Miniprep II kit (Zymo Research). For enrichment steps, high-fidelity PCR of the selected variants was carried out with Phusion™ High-Fidelity DNA Polymerase (Thermo Scientific) and used to directly transform yeasts; for subsequent mutagenesis cycles, the high-fidelity PCR product from each third enrichment was used as template for the next round of error-prone mutagenesis (described above). Each cycle of enrichment or mutagenesis involved transformation and collection of selected colonies. For yeast strain validation, cells were transformed with the full p415 plasmids expressing SpCas9 or CasΦ2 WT using the same screening conditions. Yeast colonies were quantified with ImageJ, manually adjusting the threshold for optimal discrimination. Each yeast experiment was performed by plating a defined amount of yeast suspension in n = 3 selective plates as technical replicates.

### Lentiviral vector library construction

The CasΦ2 sgRNA targeting the EGFP out of frame reporter cell line was PCR-amplified from the pUC19 plasmid prior HDV addition and sgRNA optimization and inserted into the LentiCRISPR-V1 vector using a double digestion with NdeI and EcoRI. The construct was subsequently modified to include an intron positioned upstream of the CasΦ2 cloning site, in order to prevent unwanted basal expression of the CasΦ2 variants in bacteria and avoid the possible loss of clones, as previously described [18]; the intron was PCR-amplified and inserted in LentiCRISPR-V1 CasΦ2-OOF-sgRNA by double digestion with BamHI and NheI. The resulting plasmid served as the source of the KpnI/NheI-digested backbone for preparing the lentiviral library. CasΦ2 variants from the third enrichment of the fourth round of yeast screening were amplified from plasmid DNA, digested with KpnI and NheI, and ligated into the prepared backbone at room temperature for 2 hours. The ligation mixture was purified AMPure XP Beads (Beckman Coulter) and electroporated into ElectroMAX DH5α-E Competent Cells (Invitrogen). Bacterial colonies were collected until a minimum 100x library coverage was achieved; based on the total colonies obtained from the yeast screening, the number of distinct variants was estimated at 5331. The pooled plasmid library was purified with the NucleoBond PC 500 Maxi kit (Macherey–Nagel). The primers used for lentiviral vector cloning are listed in **Supplementary Table S2**.

### Cell transduction and sorting

For the production of the lentiviral library, 2 × 10⁷ HEK293T cells (American Type Culture Collection) were transfected with 25 μg of the LentiCRISPR-sgRNA-OOF-intron-CasΦ2 library, along with 16.2 μg of pCMV-deltaR8.91 and 8.7 μg of VSV-G, using the polyethylenimine (PEI) transfection approach. The resulting lentiviral particles were passed through a 0.45-μm PES filter and concentrated by ultracentrifugation; the viral preparation was then resuspended in Opti-MEM (Gibco) and stored at −80 °C. The MOI was determined by exposing a counted number of cells to increasing amounts of viral particles and assessing their survival in puromycin (1 μg/ml) after 2 days using an MTT assay. For cell sorting, EGFP OOF 293 cells were plated and transduced the following day with the lentiviral library at an MOI of approximately 0.3, ensuring more than 100× coverage of the variant pool. Two days later, cells were placed under puromycin selection (1 μg/ml), and after additional 3 days they were sorted based on positive EGFP fluorescence using FACS ARIA III (BD Biosciences). After sorting, cells were expanded and harvested for genomic DNA extraction using the DNeasy Blood and Tissue kit (Qiagen).

### Nanopore library preparation

For UMI-tagging, CasΦ2 variants were amplified from 1 μg of genomic DNA of sorted cells using Phusion™ High-Fidelity DNA Polymerase (Thermo Scientific): 2 cycles of PCR were performed with primers carrying a randomized sequence of 20 bp (UMIs) and external tails following manufacturer’s conditions. The PCR product was purified using AMPure XP Beads (Beckman Coulter) and used as a template for 32 additional cycles of high-fidelity PCR to amplify the UMI-tagged molecules using primers annealing on the external sequence. Primers are listed in **Supplementary Table S3**. The final product was purified using AMPure XP beads and quantified by Qubit dsDNA High Sensitivity Assay kit (Invitrogen).

Sequencing libraries were prepared using the Oxford Nanopore Technologies (ONT) Ligation Sequencing Kit XL (SQK-LSK114-XL), following the manufacturer’s protocol for native, PCR-free DNA. Briefly, 280 ng of purified amplicons pool DNA was subjected to end-repair and dA-tailing using the NEBNext Ultra II End Repair/dA-Tailing Module (New England Biolabs). After bead purification, adapters were ligated using Quick T4 DNA Ligase (New England Biolabs). The ligated library was purified with ONT Short Fragment Buffer and eluted in Elution Buffer (EB) to obtain the final sequencing-ready preparation.

### Nanopore sequencing

The prepared library was loaded onto a R10.4.1 flow cell (FLO-MIN114) and sequenced on a MinION Mk1B device using MinKNOW (version 24.02.8). Active channel selection and reserved pores were enabled to maximize sequencing efficiency, while the instrument performed a pore scan every 1.5 hours. Real time basecalling was performed only to monitor the run using the Fast model at 400 bps, with a minimum read length threshold of 200 bp and a minimum Q score of 8; modified basecalling was not enabled. Data output was configured such that FASTQ files were generated every 10 minutes, while POD5 files were generated on an hourly basis; FAST5 output was disabled. Sequencing was carried out for 72 hours, and the generated raw files were collected for downstream analysis. The generated FASTQ files were discarded and a new accurate basecalling was performed from POD5 files.

### Nanopore data analysis

Basecalling was performed using dorado v0.8.3 with parameters --min-qscore 8 -- emit-fastq --no-trim and the r1041_e82_400bps_hac_v5.0.0 model. Raw sequencing reads were trimmed using Porechop v0.2.4 with parameters --min_split_read_size 2200 --adapter_threshold 80 --min_trim_size 20 --middle_threshold 75 -- extra_end_trim 0 --extra_middle_trim_good_side 0 --extra_middle_trim_bad_side 0 -- check_reads 100000. The adaptors.py file in Porechop was modified to include customized concatenated primers originating from end-to-end ligation of PCR products. Trimmed reads were filtered using Filtlong v0.2.1 with parameters -- min_length 2200 --min_mean_q 70 and cutadapt v4.9 with parameters -m 2200 -M 2800. UMI extraction and consensus read generation were performed using the longread_umi pipeline [21] v0.3.2 (https://github.com/SorenKarst/longread_umi) with UMI patterns N_4_YRN_4_YRN_4_YRN_4_ (forward) and N_4_RYN_4_RYN_4_RYN_4_ (reverse) and customized flanking sequences. A total of 56,957 unique UMIs were identified, corresponding to 1,341,916 raw reads.

UMI-corrected consensus reads were aligned to the CasPhi2 wild-type sequence using minimap2 v2.28-r1209 with parameters -a -x map-ont. The resulting alignment was filtered with samtools v1.21 with parameters -F 2304 and processed with a customized python script. Briefly, consensus reads resulting from UMI bins containing less than 15 raw reads (cutoff for Q40 average quality in [21]) were discarded to retain only high quality consensus reads (44,874 UMI-corrected reads from 1,231,371 raw reads). Filtered reads were used to compute the mutation pileup along the protein length. Then, combinations of mutations (haplotypes) in each read were extracted and converted to amino acid substitutions, resulting in a total of 4,130 unique variants.

To generate a two-dimensional visualization of identified variants, they were first filtered to remove variants reconstructed only once, retaining a total of 2,382. Mutations were converted to one-hot encodings and those with very low variance (<5e-4) were removed, retaining 580 mutations. Variants were then clustered using K-means clustering (K=20) and visualized using UMAP. Consensus mutations were identified for each cluster as mutations present in at least half of the cluster members. Mutations were visualized in the CasPhi2 ternary complex (PDB 7LYS) using ChimeraX [24].

### In vitro PAM determination assay

The *in vitro* PAM determination of CasΦ2 was performed by adapting the protocol from Karvelis et al. [25]. CasΦ2 WT and evoCasΦ2 human codon optimized DNA sequence were cloned into the expression vector pT7-N-His-GST (Thermo Fisher Scientific) for *in vitro* transcription and translation (IVT) as described above. The IVT reaction was performed using the 1-Step Human High-Yield Mini IVT Kit (Thermo Fisher Scientific) according to the manufacturer’s protocol. A DNA fragment containing T7 promoter and CasΦ2 sgRNA sequence was synthesized as complementary oligonucleotides annealed and then *in vitro* transcribed using the HighYield T7 RNA Synthesis Kit (Jena Bioscience), following the manufacturer’s guidelines. Purification of the *in vitro* transcription reaction was done by overnight ammonium acetate precipitation. The CasΦ2-sgRNA ribonucleoprotein (RNP) complex was assembled by combining 20 μL of the supernatant containing soluble CasΦ2 protein with 1 μL of RiboLock RNase Inhibitor (Thermo Fisher Scientific) and 2 μg of sgRNA. The RNP was incubated at room temperature for 30’. The RNP was subsequently incubated with 1 μg of 7N randomized PAM plasmid DNA library [6] and cleavage buffer (10 mM Hepes-K pH 7.5 RT, 150 mM KCl, 5 mM MgCl2, 0.5 mM TCEP) [11] at 37 °C for 3 h. The ends of the cleaved PAM library were repaired with 1 μL T4 DNA polymerase and dNTPs, 3’-dA overhangs were added with 1 μL DreamTaq, and RNA was removed with RNase A/T1. The final product was purified using a GeneJet PCR Purification Kit (Thermo Fisher Scientific). A double-stranded DNA adapter was ligated to the PAM library DNA ends, and the final ligation product was purified with AMPure XP beads (Beckman Coulter), with a 1:0.8 ratio. A first PCR (Phusion HF DNA polymerase, Thermo Fisher Scientific) was performed to enrich the cleaved sequences using a forward primer annealing on the adapter and a reverse primer designed on the plasmid backbone downstream of the PAM [6]. A second round of PCR was performed to attach Illumina indexes and adapters. PCR products, in both steps, were purified using purified AMPure XP beads (Beckman Coulter) (ratio 1:1). The library was analyzed by 250 bp paired-end reads sequencing, on an Illumina MiSeq sequencer. PAM sequences were extracted from Illumina MiSeq reads and used to generate PAM sequence logos, using Logomaker version 0.8 [26]. PAM heatmaps [27] were used to display PAM enrichment, computed dividing the frequency of PAM sequences derived from target cleavage events by the background frequency of uncleaved sequences.

### Indels analysis at human genomic loci

To assess genome editing efficiency, HEK293T cells were seeded in 24-well cell culture plates and, on the following day, co-transfected with 400 ng of the sgRNA expression plasmid and 650 ng of the CasΦ2 plasmid. Transfections were performed using TransIT-LT1 reagent (Mirus Bio) following the manufacturer’s instructions. Cells were selected in puromycin (1 μg/ml) after 2 days and harvested after 3 additional days. For TIDE analysis, genomic DNA was extracted using QuickExtract DNA Extraction Solution (Lucigen); the targeted genomic regions were then amplified by PCR with HOT FIREPol MultiPlex Mix (Solis BioDyne), subjected to Sanger sequencing, and the resulting data were analyzed using the TIDE software [28]. Primers used for TIDE analysis are reported in **Supplementary Table S2**. For deep amplicon sequencing, genomic DNA was extracted using the DNeasy Blood and Tissue kit (Qiagen); target regions were first amplified by an initial round of PCR carried out with Phusion™ High-Fidelity DNA Polymerase (Thermo Scientific), using 400 ng of genomic DNA as template. The resulting amplicons were subsequently purified with AMPure XP beads (Beckman Coulter), quantified by Qubit dsDNA High Sensitivity Assay kit (Invitrogen), and subjected to a second high-fidelity PCR in order to incorporate Illumina indexing adapters. Following a final purification and quantification step, equimolar amounts of each amplicon were pooled and sequenced with a V2 flow cell on a MiSeq platform (Illumina, 150 bp, paired end). Editing levels were assessed using CRISPResso2 [29] v2.3.1 with parameters -- default_min_aln_score 30 --min_bp_quality_or_N 20 --ignore_substitutions -- cleavage_offset 1 -w 20, running in mixed pooled mode and aligning reads to the GRCh38.p14 human genome assembly. Primers used for deep amplicon sequencing are listed in **Supplementary Table S3**.

## DATA AVAILABILITY

Nanopore sequencing data for the CasΦ2 variant library and deep sequencing for genome editing quantification is available on NCBI (BioProject PRJNA1445222). The code used in this study to analyze the nanopore library is available on GitHub: https://github.com/Matteo-Ciciani/EPICA-2.

## ACKNOWLEDGEMENTS

We are grateful to Cereseto’s lab for helpful discussion throughout the project. This work was supported by the European Union’s Horizon Europe EIC Pathfinder Programme under grant agreement n. 101071041 (AAVolution) and by the Italian Ministry of University and Research (MUR) under the Fondo Italiano per la Scienza FIS program project “Microbiome yielded Biotechnological Evolution-based Therapies” grant n. FIS00002542.

## AUTHOR CONTRIBUTIONS STATEMENT

G.V.R, M.D.G. designed and performed the experiments; M.C. developed the computational pipeline for sequencing analyses; V.D.S, R.B., C.V. designed and performed the long-read nanopore sequencing experiments. G.V.R, M.D.G., D.M., E.K., A.Co collected and analyzed the data. A.Ce., A.Ca, G.V.R, M.C. conceived and designed the study, wrote and edited the paper; A.Ce. was responsible for the coordination of the study. All authors read, corrected, and approved the final manuscript.

