## Supplementary figures for "Enhancing hypercompact CasΦ2 activity through EPICA.2, an optimized eukaryotic directed evolution platform"

### Current affiliation:

1. Department CIBIO, Laboratory for Advanced Genome Editing Technologies, University of Trento (Italy)
2. Department CIBIO, Laboratory of Computational Metagenomics, University of Trento (Italy)
3. NGS Core facility, University of Trento (Italy)
4. Alia Therapeutics s.r.l, Trento (Italy)

### Anemocyte, Via R. Lepetit 34, Gerenzano, Italy

#### Supplementary Figure 1

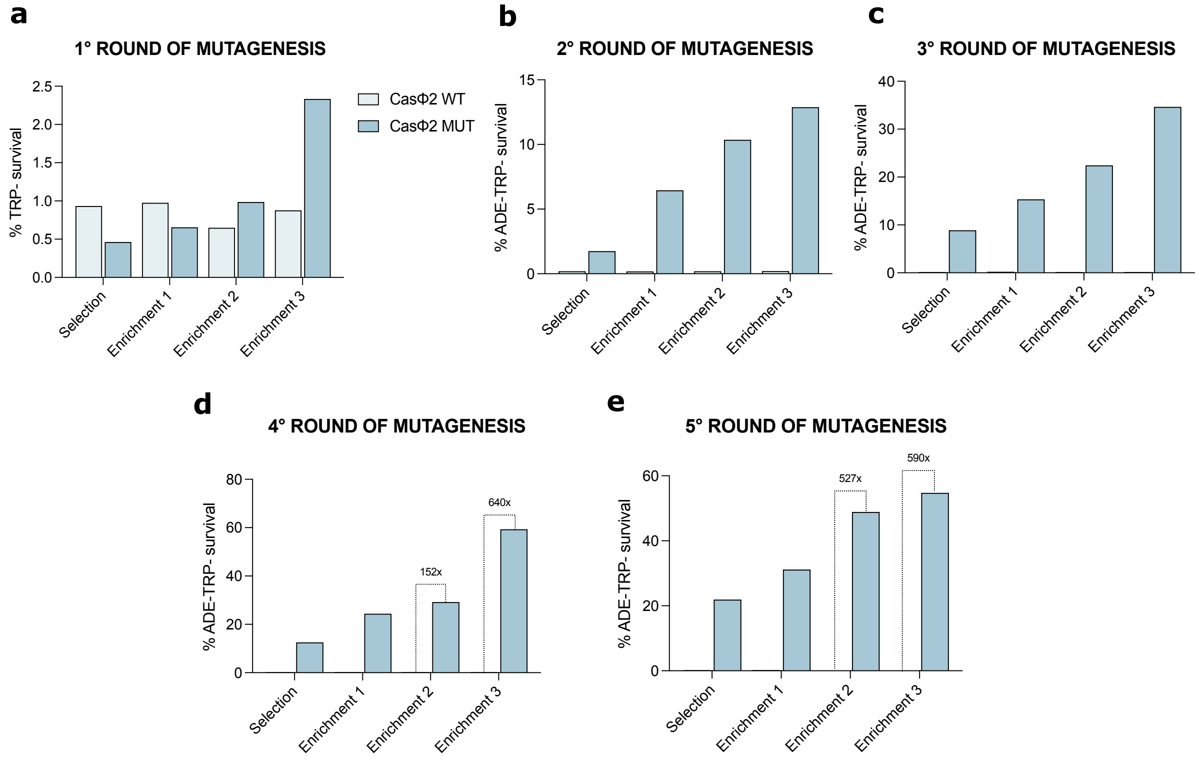


**Supplementary figure 1. Yeast screening for CasΦ2 activity enhancement.** (**a**) Activity of the evolved CasΦ2 mutant variants (CasΦ2 MUT) from the first round of mutagenesis in comparison with CasΦ2 WT at each enrichment. Survival percentages are expressed as the ratio of colonies growing on TRP− plates to the total number of transformants obtained under non-selective conditions. (**b**) Activity of the evolved CasΦ2 mutant variants (CasΦ2 MUT) from the second round of mutagenesis in comparison with CasΦ2 WT at each enrichment. Survival percentages are expressed as the ratio of colonies growing on TRP−ADE- plates to the total number of transformants obtained under non-selective conditions. (**c**) Activity of the evolved CasΦ2 mutant variants (CasΦ2 MUT) from the third round of mutagenesis in comparison with CasΦ2 WT at each enrichment. (**d**) Activity of the evolved CasΦ2 mutant variants (CasΦ2 MUT) from the fourth round of mutagenesis in comparison with CasΦ2 WT at each enrichment. (**e**) Activity of the evolved CasΦ2 mutant variants (CasΦ2 MUT) from the fifth round of mutagenesis in comparison with CasΦ2 WT at each enrichment.

**Supplementary Figure 2**

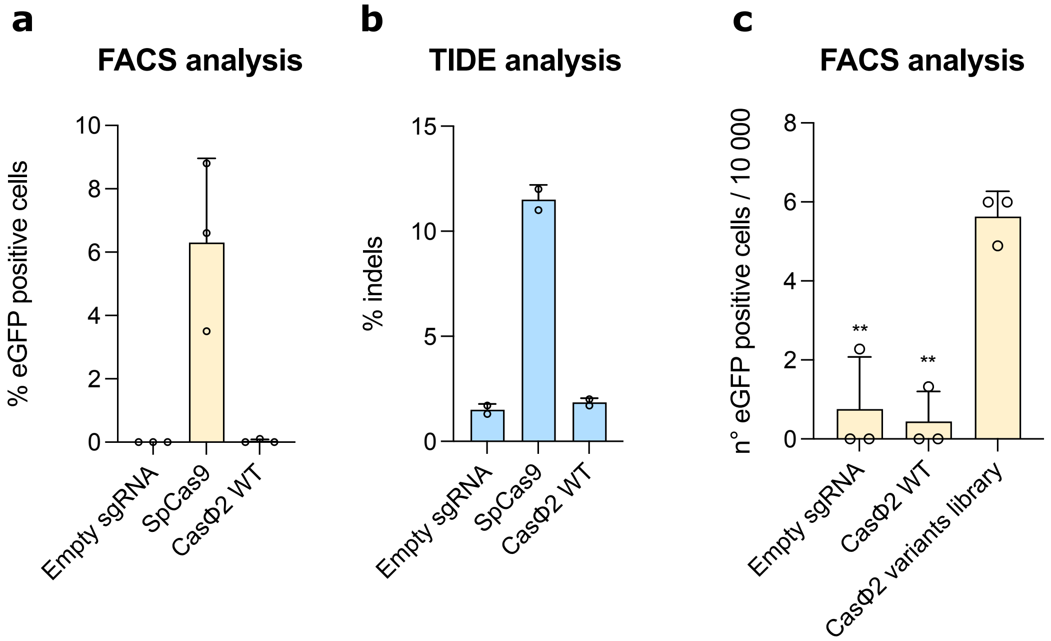


**Supplementary figure 2. Screening of enhanced CasΦ2 variants in mammalian cells.** (**a**) Validation of the Cas-mediated EGFP frameshift reporter cell line with SpCas9 and CasΦ2 WT. Percentages of EGFP positive cells obtained by FACS analysis 5 days after transduction with lentiviral vectors carrying SpCas9 and CasΦ2 WT with sgRNAs targeting EGFP out of frame reporter. (**b**) Editing efficiency of SpCas9 and CasΦ2 WT on the target site of the Cas-mediated EGFP frameshift reporter cell line. Percentages of indels are obtained by TIDE analysis 5 days after transduction with lentiviral vectors carrying SpCas9 and CasΦ2 WT with sgRNAs targeting EGFP out of frame reporter. (**c**) Percentages of EGFP positive cells obtained by FACS analysis 5 days after transduction with lentiviral vectors carrying CasΦ2 WT and CasΦ2 variants from the third enrichment of the fourth round of mutagenesis of yeast screening using sgRNAs targeting EGFP out of frame reporter. In **a**, **b** and **c** data are reported as mean ± standard deviation of $n \geq 2$ biologically independent samples. Individual values are represented as empty circles. Statistical analysis in **c** was performed using unpaired t test: ** $P < 0.01$, *** $P < 0.001$.

#### Supplementary Figure 3

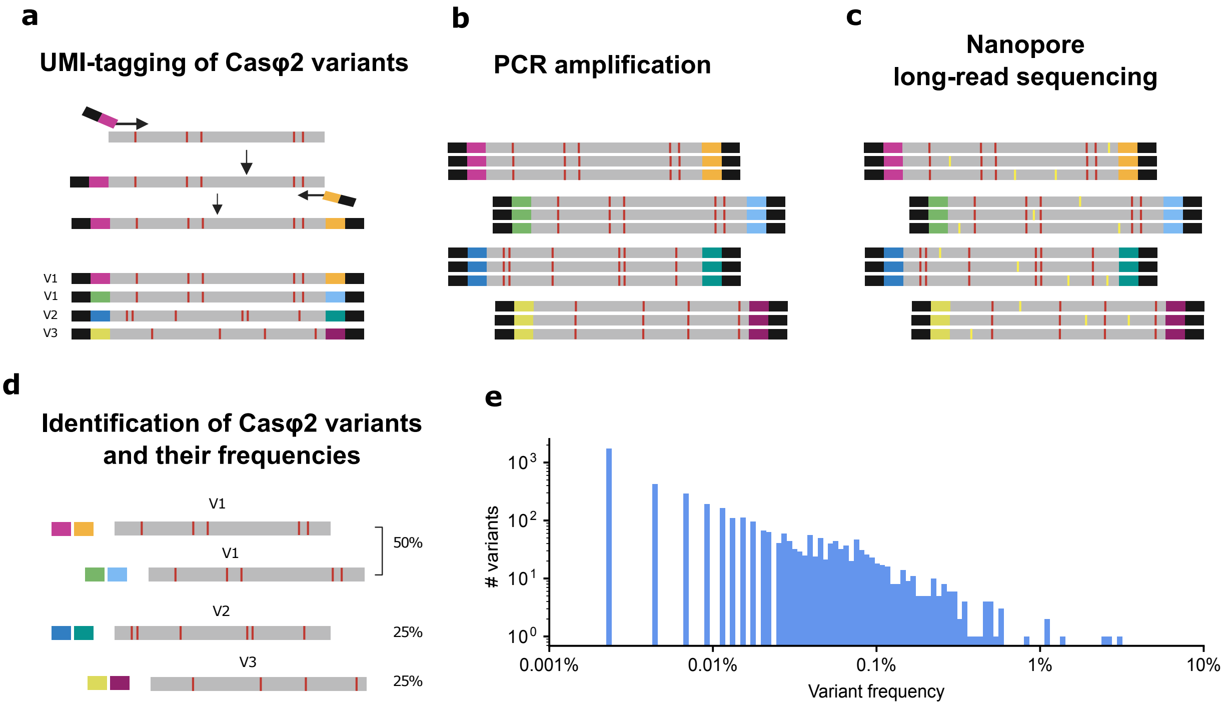


**Supplementary figure 3. Long-read sequencing of CasΦ2 variants screened in mammalian cells.** (**a**) CasΦ2 variants with mutations (red barlines) from the previously sorted cells are amplified with the addition of UMI sequences (UMI-tagging) through 2 cycles of PCR. (**b**) Multiple rounds of PCR are performed on UMI-tagged CasΦ2 variants using external primers (black) to produce multiple copies of each mutant. (**c**) CasΦ2 variants copies are long-read sequenced though Oxford Nanopore Technologies; sequencing errors (yellow barlines) are distributed across sequences of CasΦ2 variants multiple copies while mutations (red barlines) are maintained in different copies of each variant. (**d**) CasΦ2 variants are identified through error correction which determines effective mutations (red barlines); variants presenting the same mutations which are tagged with different UMIs are then grouped together to establish frequencies of the mutant in the library. (**e**) Frequency distribution of CasΦ2 variants after long-read sequencing.

#### Supplementary Figure 4

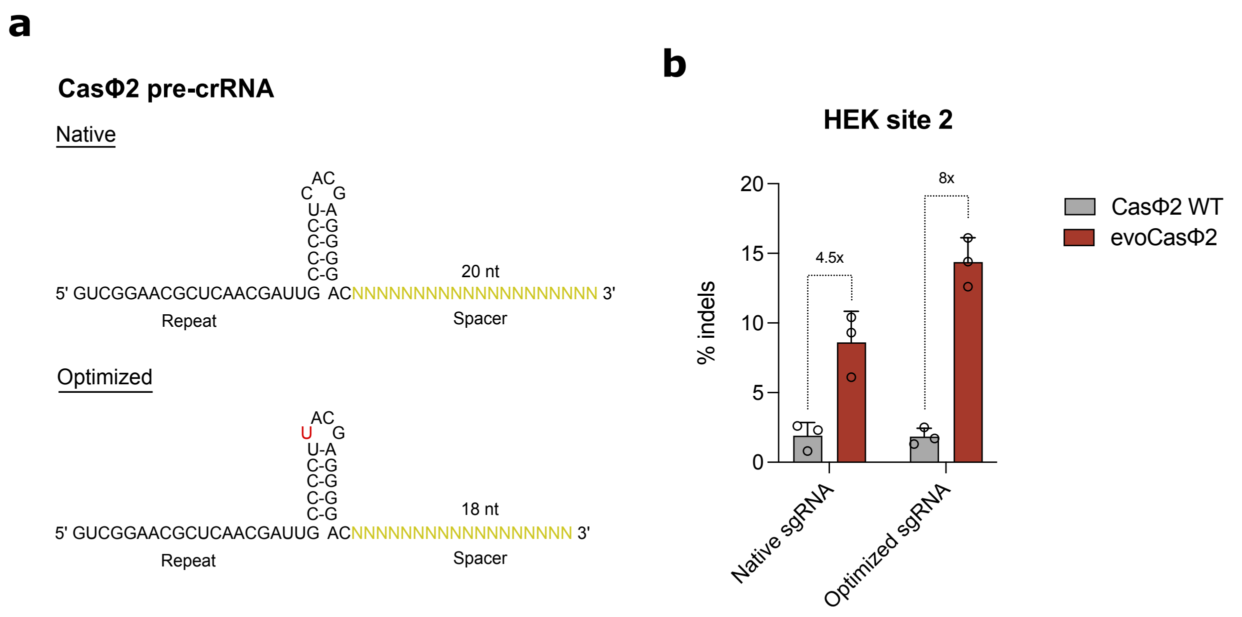


**Supplementary figure 4. Optimization of CasΦ2 sgRNA architecture.** (**a**) Representation of modifications introduced in the CasΦ2 sgRNA to enhance activity. One of the cytidines of the pre-crRNA stem loop is converted in a U (red) and the spacer (yellow) length is reduced from 20 nt to 18 nt. (**b**) Editing efficiency of CasΦ2 WT and evoCasΦ2 in HEK site 2 with native and optimized sgRNA. Percentages of indels are obtained by TIDE analysis 5 days after transfection of sgRNA and Cas plasmids in HEK 293T cells. In **b** data are reported as mean ± standard deviation of n = 3 biologically independent samples. Individual values are represented as empty circles.

#### Supplementary Figure 5

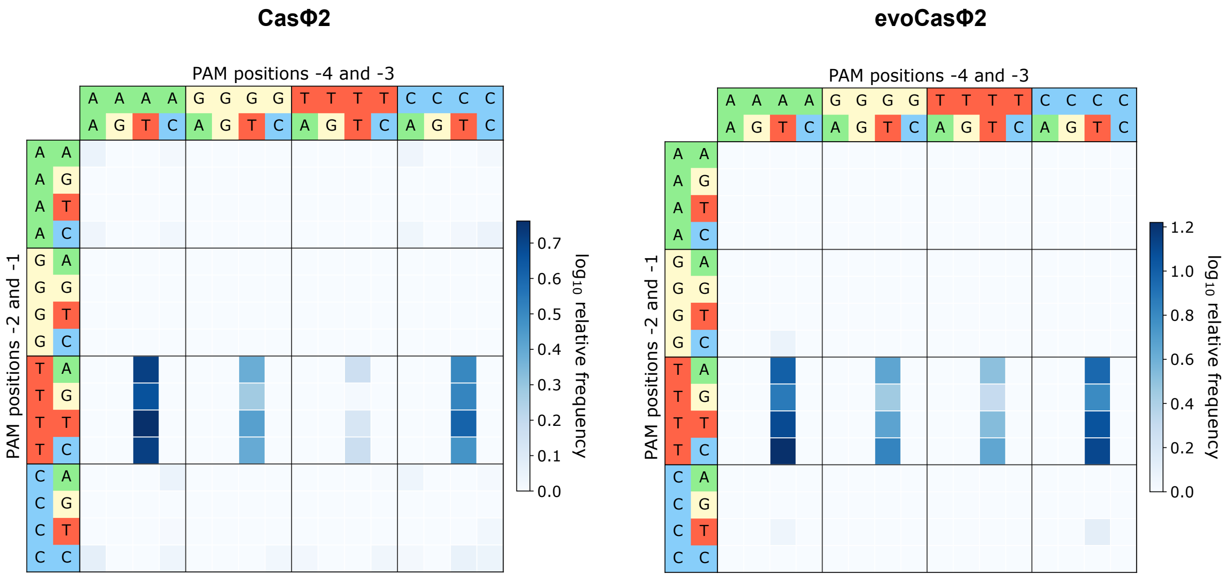


**Supplementary figure 5. *In vitro* PAM characterization of CasΦ2 WT and evoCasΦ2.** Heatmaps representation of CasΦ2 WT and evoCasΦ2 PAM preference obtained by *in vitro* determination PAM assay. Consensus sequences of PAM recognitions are illustrated in **Figure 3c**.
